# Most early-born subplate neurons persist as Layer 6b neurons in the adult mouse neocortex

**DOI:** 10.1101/2025.11.20.689634

**Authors:** Yusuke Sugita, Keiko Moriya-Ito, Carina Hanashima, Chiaki Ohtaka-Maruyama

**Affiliations:** Developmental Neuroscience Project, Department of Brain & Neurosciences, Tokyo Metropolitan Institute of Medical Science, Tokyo, Japan; Graduate School of Advanced Science and Engineering, Waseda University, Tokyo, Japan

**Keywords:** Neocortex, Subplate, postnatal development, birthdate analysis, tissue-clearing

## Abstract

Subplate neurons (SpNs) are among the earliest-born neurons in the mammalian neocortex and play key roles in radial migration and transient circuit formation. It has long been assumed that most SpNs undergo extensive postnatal cell death, leaving only a small remnant population that contributes to layer 6b (L6b) in adulthood. However, the extent to which SpNs actually persist as L6b neurons has remained unresolved, partly because previous studies lacked quantitative, whole-cortex analyses that account for postnatal cortical expansion. Here, we performed a comprehensive birthdating analysis using multiple EdU injections spanning the entire neurogenic window of SpNs in mice, combined with whole-neocortex 3D tissue clearing to measure subplate and L6b volumes. This approach allowed us to directly estimate the total number and distribution of SpN-derived neurons in adulthood. We found that most early-born SpNs persist as L6b neurons, and that the apparent postnatal reduction in SpN density reflects tangential cortical expansion rather than neuronal loss. Moreover, surviving SpNs comprise diverse neuronal subtypes reminiscent of those present at early postnatal stages. Together, these findings demonstrate that, in rodents, the majority of adult L6b neurons originate from SpNs, revising the long-held view that the subplate is a largely transient neuronal population.

## Introduction

The mammalian neocortex consists of a distinct six-layer structure, with Layer 6 b (L6b) as the thin cell layer separating the neocortex from the white matter (WM). L6b contains L6b neurons (L6bN), which project to multiple intercortical and extracortical regions, including layer 1, layer 5a, and the medial posterior thalamic nucleus (POm) (Chang et al. 2024; Hoerder-Suabedissen and Molnar 2013; Hoerder-Suabedissen et al. 2018; Kim et al. 2025; Viswanathan et al. 2017; Zolnik et al. 2024). Emerging evidence highlights the unique roles of L6b neurons in the adult brain, such as regulating sleep-wake states and cognitive functions (Ben-Simon et al. 2022; Clancy and Cauller 1999). Notably, a subset of deep-layer neurons specifically express orexin receptor type 1 and type 2 mRNA (Tsuneoka and Funato 2024), and only L6b neurons respond to Orexin B across the neocortex, implicating that SpN may be involved in the sleep state regulation (Bayer et al. 2004). Furthermore, entorhinal L6b neurons project to the hippocampus, and their pathway contributes to spatial coding and memory in mice (Ben-Simon et al. 2022). Therefore, despite their small numbers, L6b neurons play essential role in integrating diverse information in the adult neocortical circuitry. However, the developmental origin of L6b neurons remains unclear. In the developing neocortex, the deepest layer of the cortical plate, corresponding to L6b in the adult, is called the “subplate (SP)” (Allendoerfer and Shatz 1994; Feldmeyer 2023; Kanold and Luhmann 2010; Molnar et al. 2020). The SP consists of subplate neurons (SpNs) and abundant extracellular matrix (ECM). SpNs, produced during the earliest phase of neurogenesis, together with Cajal-Retzius (CR) cells are key components of the preplate (Hoerder-Suabedissen and Molnar 2015; Kanold 2009; Luhmann et al. 2018; Ohtaka-Maruyama 2020; Wang et al. 2010) and play crucial roles in forming the embryonic neocortex. One function is to form transient synaptic connections with migrating excitatory neurons, promoting their switch in migration mode (Ohtaka-Maruyama et al. 2018). Another function is the temporal establishment of thalamocortical axon (TCA) connections from the thalamus to SpNs, which subsequently guide the TCA projection to the layer 4 neurons, facilitating the formation of mature thalamocortical circuits (Doyle et al. 2021; Hoerder-Suabedissen and Molnar 2015; Meng et al. 2014; Molnar and Kwan 2024).

Although SpNs and L6b neurons are located in the same position, the deepest part of the cortical plate, they are considered distinct due to their functional differences (deFazio et al. 1987; Marx et al. 2017; Rosen and Harry 1990; Woo et al. 1991). Alternatively, study also suggest that a subset of SpNs may survive as a uniform layer, referred to as layer 6b or layer 7 (Marx et al. 2017). During this process, the decrease in SpN density is thought to result primarily from cell death in the SP in the rodent neocortex(Allendoerfer and Shatz 1994). Earlier birthdating studies in rats suggested that a substantial fraction of SpNs may survive into adulthood and contribute to L6b (Valverde et al. 1995). However, these analyses relied on single-pulse radiolabeling that captures only a very narrow birthdate window and labels only a small subset of SpNs, and they lacked quantitative, whole-cortex assessment. As a result, the persistence and abundance of SpN-derived L6b neurons have remained unresolved. This uncertainty arises from several technical challenges (Feldmeyer 2023; Marx et al. 2017). First, the molecular identities of SpNs are heterogeneous, and the absence of universal SpN-specific markers complicates their comprehensive characterization. Second, postnatal expansion of neocortical volume, including the SP, makes it difficult to assess the precise structure and distribution of SpNs across the entire neocortex. Third, birthdate analysis using a single injection of thymidine analogs provides only a short labeling window (∼four hours), capturing only a subset of early-born SpNs(deFazio et al. 1987). Moreover, although previous studies suggested that the developmental decline in SpN density might reflect overall brain growth (Rosen and Harry 1990; Woo et al. 1991), they did not directly measure postnatal volume changes in the SP or L6b, leaving the actual change in SpN number uncertain and the long-standing assumption of postnatal SpN loss in need of re-evaluation.

To overcome these limitations, we performed extended-window EdU birthdating by repeatedly administering EdU from E10.5 to E12.5, allowing comprehensive labeling of early-born SpNs. We then applied whole-brain CUBIC tissue clearing (Matsumoto et al. 2019) to reconstruct the entire SP and L6b in three dimensions and quantify their volumes. By combining volumetric measurements with NeuN-based cell counts at P0 and 8 weeks, we estimated the total number of SpN-derived neurons across the neocortex.

These analyses revealed that, at least in rodents, most early-born SpNs persist into adulthood, contrary to the long-held assumption of extensive postnatal SpN loss. Our findings demonstrate that adult L6b neurons are mostly derived from surviving SpNs and that the early-born SpN population does not undergo significant numerical reduction during postnatal development.

## Materials and Methods

### Animals

All animal procedures were conducted in accordance with the guidelines of the Tokyo Metropolitan Institute of Medical Science Animal Care and Use Committee. All experimental protocols were approved by the Committee under the following approval numbers: 21-070, 22-079, 23-008, 24-005, and 25-014. The Lpar1-GFP males [Tg(Lpar1-EGFP) GX193Gsat/Mmucd (030171-UCD, MMRRC)] were mated with ICR female mice in order to acquire more pups than mated with B6 black female and bear the stress from the multiple injection of thymidine. Wild-type (WT) ICR females (supplied by LSC, Japan) were mated for 12 h overnight with Lpar1-GFP males and checked for plugs. E0.5 was defined as the first midday after checking plugs.

### EdU treatment

Pregnant dams were injected with 15 mg/kg EdU in sterile saline (Mase et al. 2021) on E12.5. For E11 whole labeling (E11wl), intraperitoneal injection of EdU was performed 6 times in each 4 h from E10.5 to E11.5. For E12 whole labeling (E12wl), 4 times in each 6 h from E11.5 and E12.5 during 24 h. WT and Lpar1-GFP (male, Lpar1-GFP transgene is located on the Y chromosome) pups were perfused with 4% paraformaldehyde (PFA) on E17.5, P0, P14, and 8 weeks (P56-60). The brains were dissected out and post-fixed in 4% PFA for 24 h.

### Antibodies

The following antibodies were used: anti-CTGF (goat, SANTA CRUZ, sc-14939), anti-NeuN (mouse, CST, 94403), anti-Nurr1(goat, R&DSYSTEM, AF2156), anti-GFP (chicken, Abcam, ab13970), anti-Tbr1(rabbit, Abcam, ab31940), anti-Caspase-3 (rabbit, CST, 9661S). Secondary antibodies were anti-mouse 405 (donkey, Jackson), anti-goat 488 (donkey, Invtrogen), anti-chick 488 (donkey, Jackson), anti-mouse Cy5 (donkey, Jackson), anti-rabbit Cy5 (donkey, Jackson), anti-goat 633 (donkey, Invtrogen), and anti-rat 633 (goat, Invtrogen). The antibodies were used at a 1:500 dilution.

### Immunohistochemical staining and EdU staining

The dissected brains were fixed in 4% paraformaldehyde (PFA)/PBS overnight at 4 °C. The tissue were cryoprotected in 15% sucrose/PBS overnight, followed by 30% sucrose/PBS overnight at 4 °C. The tissues were cut coronally at 20 μm on a cryostat (CryoStar, NX50) for brains aged either E17.5 and P0 sample and 30μm on a microtome (Thermo Fisher Scientific, HM440) in P4, P7, P10, P14 and 8 weeks sample.

The sections were soaked in PBS for 10 min three times and pre-incubated with 0.3% Triton X-100/PBS for 2 hr, which was then incubated overnight at 4 °C with primary antibodies diluted with PBS containing 0.5% skim milk. After washing three times with PBS, the sections were incubated with second antibodies. Then, sections were mounted with PermaFluor (Thermo Scientific) after DAPI staining (5 μg/ml, Sigma-Aldrich). EdU staining was performed by Click-iT^TM^ EdU Alexa Fluor^TM^ 555 Imaging Kit (Invitrogen by Therme Fisher Scientific, 2765704). Images of sliced sections of immunostaining were acquired by confocal microscopy (Zeiss, LSM710).

### Definition of Subplate

The subplate layer was defined as a 50 μm thick band from the bounder of WM (Hoerder-Suabedissen and Molnar 2013) on E17.5 and P0. On P14 and 8 weeks, adult L6b was also defined as 50 μm thick because we measured the width from WM by referring to the expression of CTGF and EdU.

### Tissue Clearing

We used CUBIC tissue clearing (Matsumoto et al. 2019) and applied this method to the P0 brain (Alsina et al. 2024). Lpar1-EGFP mice in P0 and 8 weeks were fixed by perfusing 4% PFA for 3 days at 4 °C. The fixed brains were washed 3 times in PBS for 6 h at room temperature (RT) and incubated in 4 mL 0.5 × CUBIC-L (1:1 N-Butyldiethanolamine and TritonX-100 in water) in PBS overnight. The solution was after that replaced with 2 mL 1× CUBIC-L and dilapidated 3 times for each overnight at 37 °C with gentle shaking. The samples were washed 3 times in PBS for 6 h at RT. The samples were incubated in 0.5 × CUBIC-R (AnTipyrine, N-Methylnicotinamide, and N-Butyldiethanolamine in water) for 6 h. Then, they were replaced twice with 1 × CUBIC-R for each overnight at 37 °C with gentle shaking. Cleared whole brain tissues were imaged by Light sheet microscopy (Olympus, MVX10-LS), and the z-axis interval was 4 μm in the P0 sample and 9 μm in the 8 weeks.

### Calculation of SP density

The SP was defined by the SpN’s marker positive (Lpar1-GFP +) area. To compare the previous studies’ definition of the SP, the area of the SP was measured. The SpNs were detected by NeuN+ in the SP and counted by ImageJ. Then, calculating the density of SpNs, the number of SpNs was divided by the area of the SP.

### Cell count

The cell count was performed by the method described below in order to avoid regional polarization of each targeted cell number in the neocortex. In each sample, two series of coronal slices were selected, which were relatively the same position as No. 54 and No. 69 in Allen Brain Reference Atlases Adult Mouse coronal Section (https://atlas.brain-map.org/atlas?atlas=602630314). In each section, three images were acquired, which are shown as black square regions in Fig S1A and S1B. Quantification was then performed on each image. In one sample, a total of six data from six images were quantified and averaged as one data.

### 3D reconstruction

The data was acquired from imaging by light sheet microscopy. On each image, the GFP+ area specified in the neocortex was cropped and the area was measured by ImageJ. These measured area data were integrated into the volume (V) with a z-axis interval of 4μm in P0 and 9 μm in 8 weeks and magnification was 2-fold at P0 and 1-fold at 8 weeks.

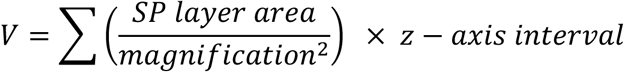

### Estimation of the number of SpNs

We calculated the density of SpN in mm^3^ by dividing thickness of each stage: 20 μm at P0 and 30 μm 8 weeks. We multiplied the density and volume of SpL to estimate the number of SpNs.

### Statistical analysis

Statistical analyses were performed using Microsoft Excel. All data were expressed as the mean ± S.D., and Welch’s t-tests (unpaired two-tailed) were used to compare the means of the two groups.

## Results

### EdU administration over a 24-hour period sufficiently capture early-born SpNs

To analyze the distribution of subplate neurons across multiple stages, we defined SpNs as early-born neurons in the neocortical neurogenesis period E10.5 to E12.5 located at the bottom edge of the cortical plate in mice (Hoerder-Suabedissen and Molnar 2013). To comprehensively label SpNs generated during this period, we employed multiple injections of ethynyl deoxyuridine (EdU), considering the efficient labeling window of a single injection of thymidine analog is 2 to 6 h (deFazio et al. 1987). To avoid EdU overdose, we divided SP neurogenesis period into two phases, E10.5 to E11.5 and E11.5 to E12.5. Six EdU injections were administered at 4-hr intervals in each phase. For the E10.5-E11.5 whole labeling (E11wl) (Fig 1A), 60% of pups survived and grew to adulthood (n= 6 pregnants). Using the same method to label SpNs generated during E11.5 to E12.5 (E12wl) resulted in almost all pups died due to stillbirth or parental abandonment (n= 12 pregnant mice). To circumvent this, we reduced EdU injections to 4 times at 6-hr intervals, which allowed the pups to survive postnatally (Fig 1D). To assess the efficiency of this protocol in labeling SpNs, we compared it to a single thymidine analog injection in previous studies. In P0 mice, the EdU+ neurons in the neocortex in mice injected with EdU at E11.5 were restricted to the SP and the marginal zone (MZ) as expected (Fig 1B), however, the density of EdU+ neurons in the SP between E11.5 single injection and E11.5 wl, the EdU+ density was approximately six times higher in E11.5 wl (Fig. 1C). Similarly, comparison of EdU+ neuron density in the SP between E12.5 single injection and E12.5 wl revealed approximately five times higher labeling in E12.5 wl (Fig 1E and 1F). These results demonstrate that multiple EdU injections at 4 or 6 hour intervals comprehensively label early-born neurons that were not achieved by a single injection. Next, we examined whether EdU+ neurons in the MZ were Cajal-Retzius (CR) cells, as CR cells are one of the earliest-born neurons in the cerebral cortex, and along with SpNs are considered to diminish during postnatal development. We tracked the localization of EdU+ SpN and CR cells during embryonic and postnatal stages. At E17.5 and P0, both EdU+ SpN and CR cells were observed in their respective locations, however, at P14, almost no EdU+ cells were detected in the MZ. In contrast, in L6b, many EdU+ neurons persisted at P14 (Fig. 1G).

**Figure 1.**
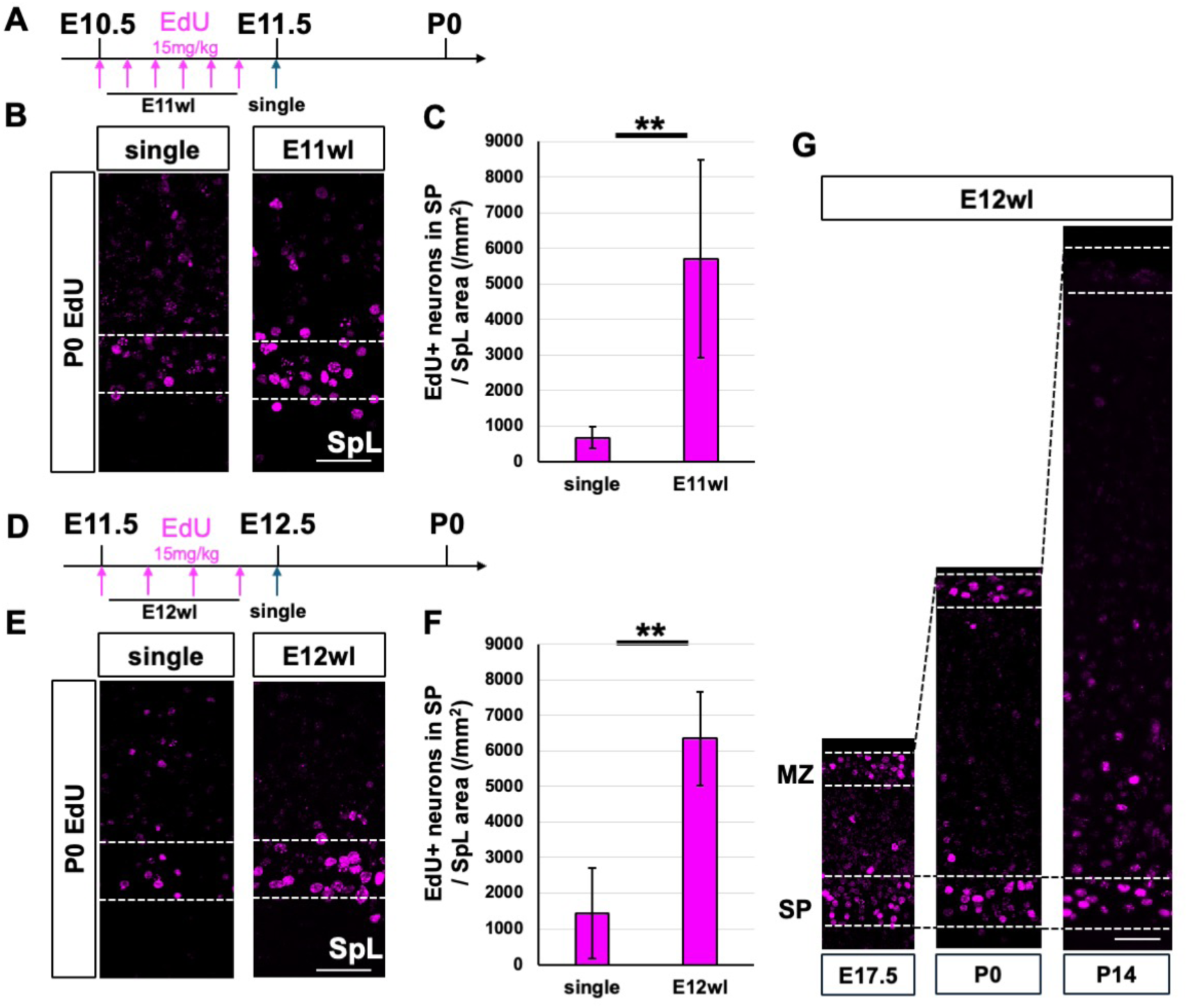
SpNs were efficiently labelled by the whole labeling method. (A) The schematic timetable of the whole labeling by EdU. For E11 whole labeling (E11wl), EdU (15 mg/kg) was injected 6 times in every 4 hours from E10.5 to E11.5. In a single-injection condition, EdU (15mg/kg) was injected once at the E11.5. Fixed samples at P0. (B) Distribution of EdU+ cells in the somatosensory cortex at P0 in the single and E11wl. Scale bars indicate 50μm. (C) The density of EdU+ neurons in the subplate layer (SpL) area in single (n = 6 field of views, 3 animals) and E12wl (n = 6 field of views, 3 animals) at P0. ^∗∗^P < 0.01; Welch’s t-tests (unpaired two-tailed) (D) The schematic timetable of the whole labeling by EdU. For E12 whole labeling (E12wl), EdU (15 mg/kg) was injected 4 times in every 6 hours from E11.5 to E12.5. In a single-injection condition, single, EdU (15mg/kg) was injected once at the E12.5. Fixed samples at P0. (E) Distribution of EdU+ cells in the somatosensory cortex at P0 in the single and E12wl. Scale bars indicate 50μm. (F) The density of EdU+ neurons in the SpL area in single (n = 6 field of views, 3 animals) and E12wl (n = 6 field of views, 3 animals) at P0. ^∗∗^P < 0.01; Welch’s t-tests (unpaired two-tailed) (G) Distribution of E12wl EdU+ cells in E17.5, P0 or P14. The upper line indicated MZ, and the bottom line indicated SP layer. Scale bars indicate 50μm.

This result suggested that EdU+ early-born neurons persisted in SP but, few in MZ at P14. These findings indicate that although SpNs and CR cells are both early-born neurons, their postnatal reduction occur in a significantly distinct manner.

### Most L6b neurons are generated between E10.5 and E12.5

The comprehensive labeling method (E11wl and E12wl) enabled us to assess the number of embryonic SP neurons that survived up adulthood as L6b neurons in the neocortex. As a result, E11wl and E12wl EdU+ cells specifically located in the L6b in the entire neocortex at 8 weeks (Fig S2 and S3). We analyzed the thickness of E12wl neurons in the entire neocortex. We measured the depth of regions in which EdU+ neurons were located in a few coronal severe sections (Fig S4 A). The thickness of EdU+ regions was about 70 μm in the anterior, about 50 μm in the somatosensory area, and 40 μm in the posterior, and the average depth in the entire EdU+ region was 50.6 μm (Fig S4B). Consistent with the notion that the SP thickness in the somatosensory cortex is 50 μm in the first postnatal week (Hoerder-Suabedissen and Molnar 2013), we identified EdU+ neurons continuously distributed in the adult neocortex in L6b within the 50 μm width from the WM. These observations indicate that SP/L6b do not expand vertically during the postnatal period.

We analyzed the ratio of EdU+ neurons in L6b neurons at 8 weeks. EdU+ early-born neurons remained in the adult L6b, and the ratio of EdU+ population at E11wl was 41.84 ±15.75 % (EdU+/NeuN+) (n= 4, Fig 2A and 2C). These results indicate that the SpNs generated through E10.5 to E11.5 represent half of the population among all SpNs, a population which survived in the adult neocortex. We further quantified E12wl EdU+ neurons, which occupied 63.03 ±11.04 % of L6b neurons (EdU+/NeuN+) (n= 4, Fig 2B and 2C). The combined results of E11wl and E12wl indicate that over 90% of L6b neurons born between E10.5 and E12.5 survive into adulthood in the neocortex (Fig 2D). In contrast, there were few EdU+/NeuN+ signals in WM, indicating that early-born neurons were strictly located inside L6b in the rodent brain. Collectively, embryonic SpNs generated during this period persists as L6b neurons and are distributed throughout the adult neocortex.

**Figure 2.**
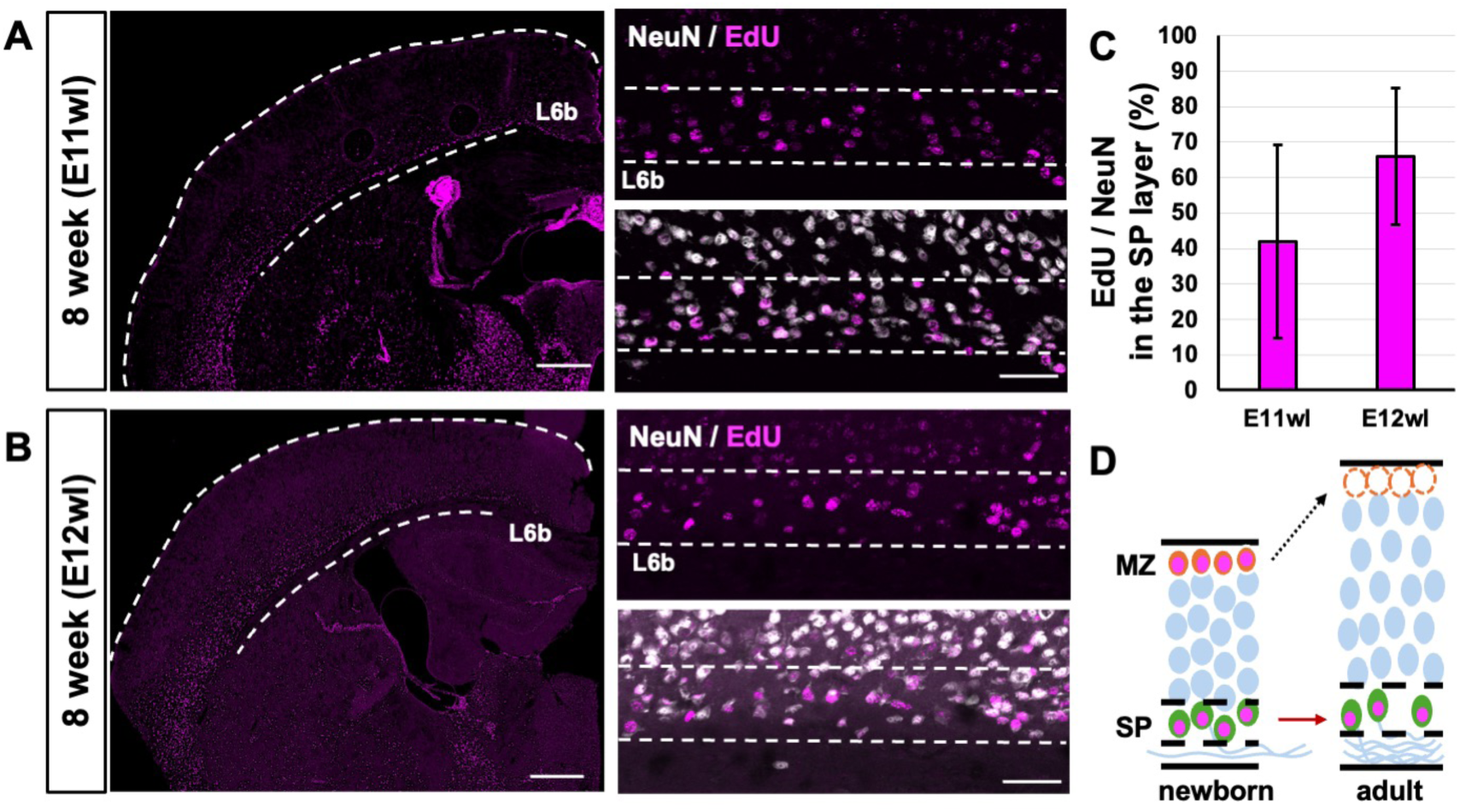
Early-born neurons persist as layer 6b (L6b) neurons in the adult cortex. (A) Confocal micrographs of E11wl (E10.5-E11.5) EdU+ cells at 8 weeks. The white dashed line indicates the lower boundary of layer 6b (L6b) in the adult neocortex. Scale bars indicate 500μm and 50μm. (B) The confocal micrographs of E12wl (E11.5-E12.5) EdU+ cells at 8 weeks. The white dashed line indicates the lower boundary of layer 6b (L6b) in the adult neocortex. Scale bars indicate 500μm and 50μm. (C) Proportion of EdU+ neurons (EdU+/NeuN+) among total L6b neurons at 8 weeks. Data are shown for E11wl at 8 weeks (n = 4 brains, n = 24 images) and E12wl at 8 weeks (n = 4 brains, n = 24 images). (D) Summary of EdU+ SpN and L6b neurons by whole labeling in postnatal development. EdU+ neurons (magenta) are located at the SP and MZ in the newborn, and no EdU+ neurons exist at the MZ, but EdU+ neurons are situated in the L6bN in the adult.

### L6b neurons have distinct molecular characters from SpNs

Previous studies have indicated the molecular diversity of SpNs during the early postnatal period (Hoerder-Suabedissen and Molnar 2013). We thus investigated whether EdU+ early-born neurons express SpN markers in the adult neocortex, and whether molecular signature between E11wl and E12wl birthdates exist. We immunostained CTGF and Nurr1 as established SpN markers, and Tbr1 as a L6 marker for E11wl and E12wl L6b (Fig 3A).

**Figure 3.**
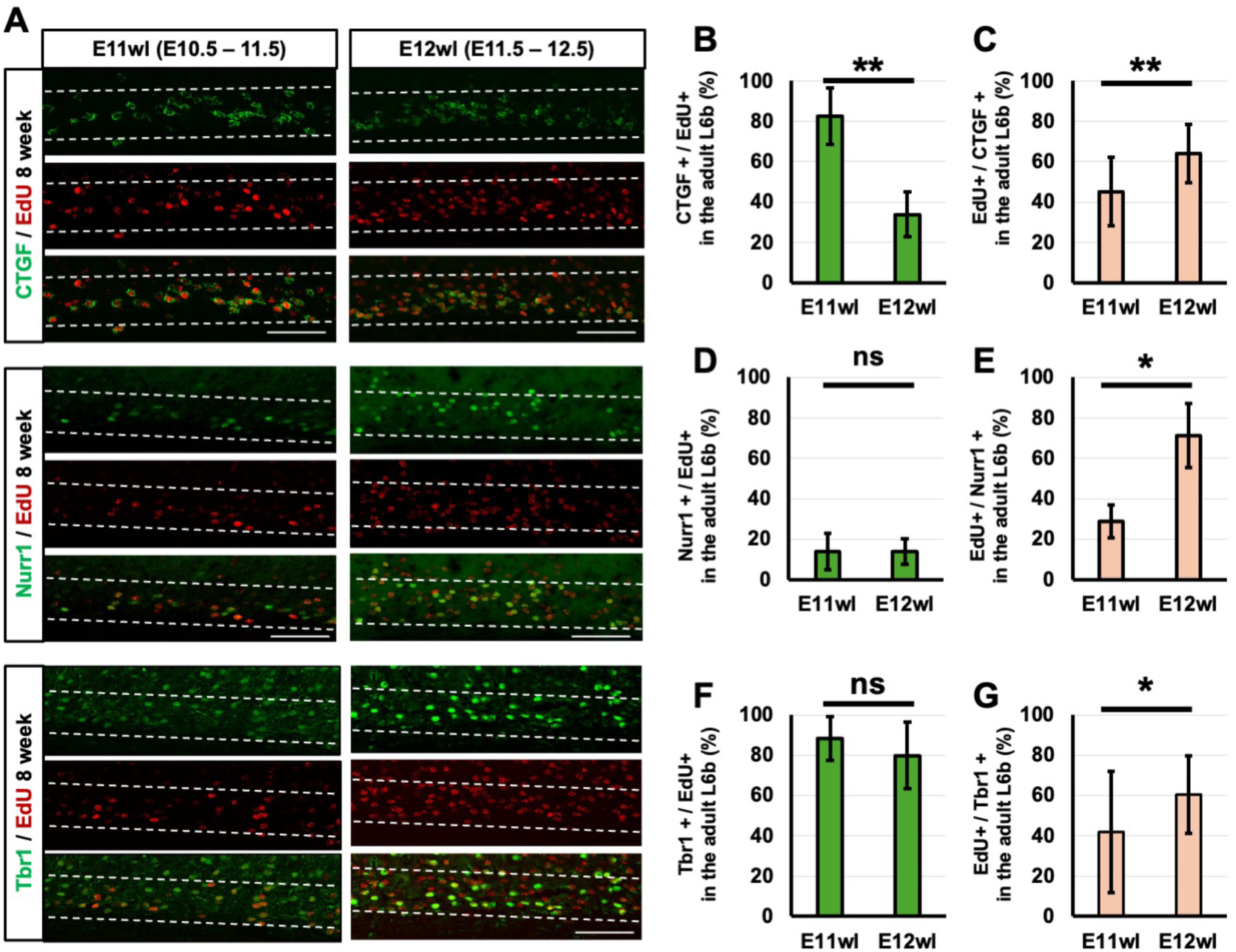
SpN marker+ neurons labeled by EdU persist in the adult L6b. (A) Immunohistochemical images showing CTGF (top), Nurr1 (middle), and Tbr1 (bottom) expression in mice subjected to E11 whole labeling (E11 WL; left) or E12 whole labeling (E12 WL; right) and analyzed at 8 weeks of age. Scale bar = 50 μm (applies to all panels). (B, D, F) Bar graphs (mean ± SD) showing the percentage of EdU+ neurons that co-expressed each marker: CTGF (B), Nurr1 (D), or Tbr1 (F) in 8 weeks (n = 4 brains, n = 24 images). ^∗^P < 0.05, ^∗∗^P < 0.01, no significant (ns); Welch’s t-tests (unpaired two-tailed) (C, E, G) Bar graphs (mean ± SD) showing the percentage of each marker+; CTGF (C), Nurr1 (E), or Tbr1 (G) neurons that colocalize with EdU+ nuclei for injections as E11wl and E12wl in 8 weeks (n = 4 brains, n = 24 images).^∗^P < 0.05, ^∗∗^P < 0.01, no significant (ns); Welch’s t-tests (unpaired two-tailed)

For SpN markers, CTGF was expressed higher in E11wl than in E12wl population (CTGF+ & EdU+ / EdU+, 82.58 ±14.18% in E11wl, and 33.92 ±11.05% in E12wl, Fig 3B). Assessing the birthdate of CTGF expressing cells within L6b neurons revealed higher E12wl population compared to E11wl (CTGF+ & EdU+ / CTGF+, 45.17 ±17.00% in E11wl, 63.95 ±14.56% in E12wl, Fig 3C). Nurr1 was also expressed at a similar ratio, however, the ratio was relatively low compared to CTGF (Nurr1+ & EdU+ / EdU+, 13.95 ±9.12% in E11wl, 13.95 ±6.43% in E12wl, Fig 3D). Notably, the Nurr1+ cell ratio was significantly different between the two birthdates, where Nurr1 represented a significantly higher number in E12wl cells (Nurr1+ & EdU+ / Nurr1+, 28.65 ±8.19% in E11wl, 71.24 ±15.98% in E12wl, Fig 3E). For both SpN markers, the total ratio in E11wl and E12wl was about 100%. Moreover, according to our result (Fig. 2C), the difference between the expression ratios was reasonable for the amount of each L6b neuron. However, the ratio of Nurr1+ in E12wl was two-fold higher than E11wl, indicating that SpNs generated at later phase of SpN-genesis are more likely to express Nurr1.

We next examined L6 marker Tbr1, which was highly expressed in both populations (Tbr1+ & EdU+ / EdU+. 88.18 ±10.75% in E11wl, 79.75 ±16.48 % in E12wl, Fig 3F). Although the ratio of Tbr1-expressing L6b neurons did not show significant differences between the two birthdates, the E12wl population represented a higher proportion of Tbr1+ cells compared to E11wl (EdU+ & Tbr1+ / Tbr1, 42.04 ±30.04% in E11wl, 60.70 ±19.16% in E12wl, Fig 3G).

Therefore, similar to Nurr1, SpNs from the later phase of neurogenesis tend to express Tbr1. Together, these results indicate that the surviving SpNs in L6b share common molecular identities with newborn SpNs. The surviving SpNs expressed both SP markers and, to some extent L6 makers, suggesting that these early-born SpNs retained in L6b may contribute to the adult neuronal circuit through the expression of these markers.

### Programmed cell death is not the main cause of the decrease in SpN density

In the adult neocortex, L6b neurons mainly consists of remnant SpNs. However, anatomical studies have shown that the density of SpNs / L6b neurons decreases during postnatal development (Hoerder-Suabedissen and Molnar 2013; Price et al. 1997; Woo et al. 1991). To confirm this, we examined the developmental changes in neuron density using NeuN immunohistochemistry as a neuronal marker. We found a reduction in the density of SP / L6b neurons during the first postnatal week (Fig S5A). It has been hypothesized that this decrease may result from programmed cell death (PCD) during postnatal development (Luhmann et al. 2018; Wong and Marin 2019). To investigate whether PCD occurs within the SP during this period, we performed immunostaining using Caspase-3 and assessed PCD at multiple postnatal stages. We first examined P4 (Fig 4A) brains due to the large decrease in cell density at this stage. Capase-3+ cells were observed throughout the neocortex but was not specifically concentrated in the SP. Immunohistochemistry at P4, P7, P10, and P14 (Fig 4B) revealed Caspase-3+ cells in the neocortex, including the SP, but the overall number of Caspase-3+ cells in the neocortex was very low (1 - 2 cells per frame of view). Moreover, comparing the number of caspase-3+ cells between the SP and L6a revealed no differences (data not shown).

**Figure 4.**
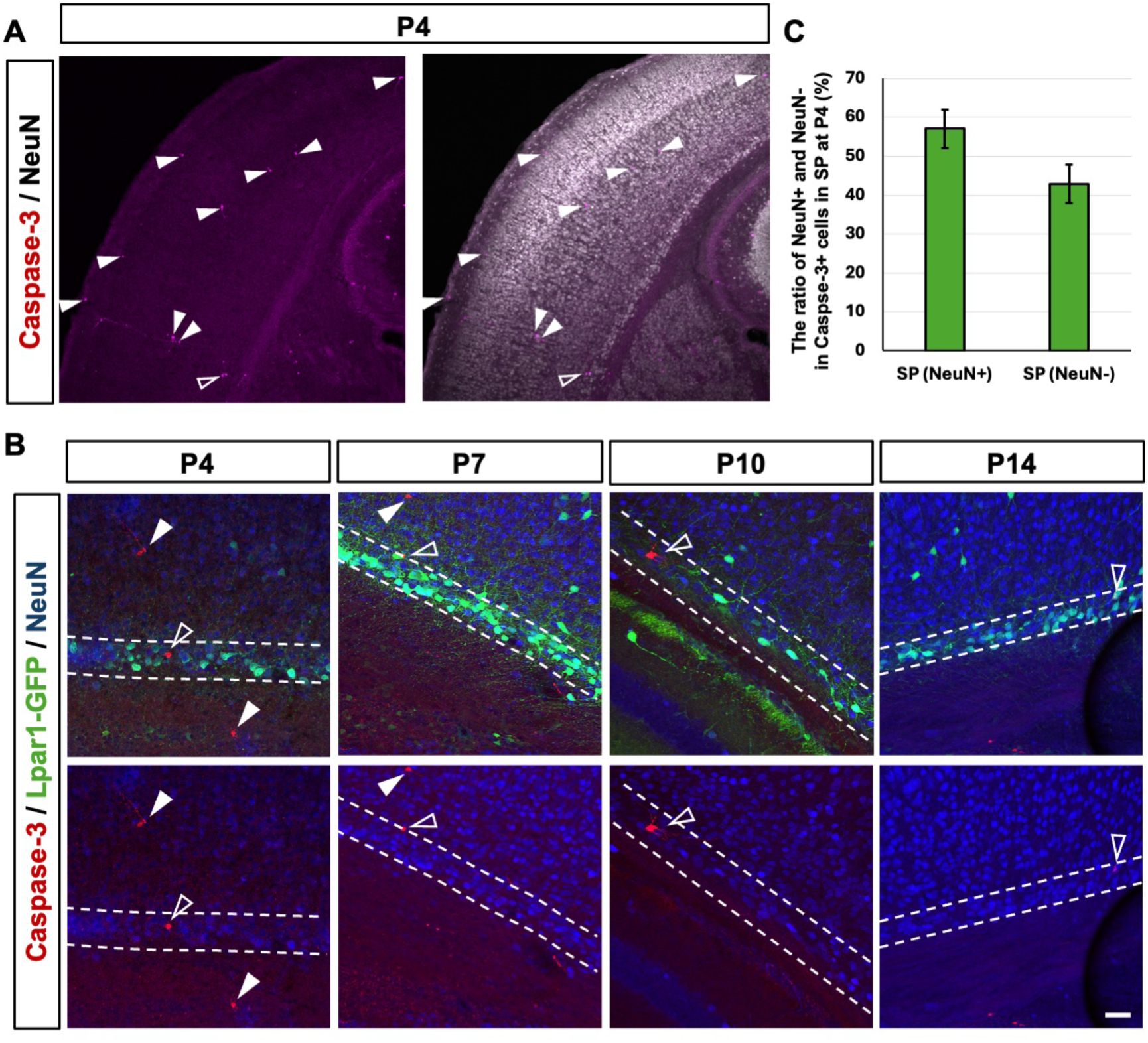
Few apoptotic neurons detected by Caspase-3 were observed in the SP during postnatal development. (A) Immunohistochemical images of Caspase-3 (magenta) and NeuN (gray) at P4. White filled arrowheads show Caspase-3+ cells in cortex and white outlined arrowheads show Caspase-3+ in SP. Scale bar = 150μm (B) Immunohistochemical images of Caspase-3 (red), Lpar1-GFP (green), and NeuN (blue) at P4, 7, 10, and 14. White filled arrowheads show Caspase-3 positive cells in cortex and white outlined arrowheads show Caspase-3+ in SP. White dashed lines denote the SP boundaries. Scale bar = 50μm (C) The percentage of NeuN+ or NeuN-among Caspase-3 positive cells in the SP at P4. Caspase-3 positive cells were counted from 8 slices in each brain and averaged as one sample.

We further assessed whether these Caspase-3+ cells within the SP include glial cells, in addition to neurons using NeuN immunostaining (Fig 4C). About 60% of Caspase-3+ cells in the SP co-expressed NeuN, confirming their neuronal identity. However, about 40% of Caspase-3+ cells did not express NeuN, indicating that either they are glial cells or apoptosis or specific cellular conditions modulate NeuN expression to circumvent their detection within Caspase-3+ cells. These findings suggest that PCD, as evidenced by Caspase-3 expression, plays a limited role in the reduction of SpN populations during postnatal development.

### Whole-brain imaging reveals minimal elimination of SpNs

The observed decrease in SpN density may be influenced by technical limitations, in which we and others calculated the total number using a series of brain sections (Fig S5A). Since these images compared different stages with same size of frame of view, without accounting for the volumetric expansion of the neocortex. Indeed, we demonstrated that both SP and L6b area expand tangentially during the first postnatal week (Fig S5B and S4C), particularly along the anteroposterior (AP)-axis of the neocortex (Fig S5D and S5E). Based on this, we hypothesized that neocortical expansion during this period may account for the decrease in SpN density.

Although previous studies attempted to account for these volume changes (Torres-Reveron and Friedlander 2007; Woo et al. 1991), their estimations were not specific to SP / L6b. To address this, we visualized the entire distribution of SpNs in the whole neocortex using the CUBIC tissue-clearing method and light-sheet microscopy at P0 and 8 weeks, and quantified the volume of SP / L6b (Matsumoto et al. 2019). We used the Lpar1-EGFP transgenic mouse line (Hoerder-Suabedissen and Molnar 2013; Walker et al. 2016) to define the SP, which consistently expresses GFP in the neurons located at SP / L6b and can distinguish the boundary between L6a and SP/L6b during postnatal development (Walker et al. 2016). We defined the SP in P0 and L6b at 8 weeks by GFP signals and measured the areas of each slice to calculate the volume of the whole SP and L6b from z-stack compilations (Fig 5A). By calculating each volume, the SP area was 0.57 ±0.04 mm^3^ (n = 4), L6b was 4.21 ±0.15 mm^3^ (n = 4, Fig 3B), and the adult L6b exhibited 7.4 fold increase in volume compared to the SP at P0.

**Figure 5.**
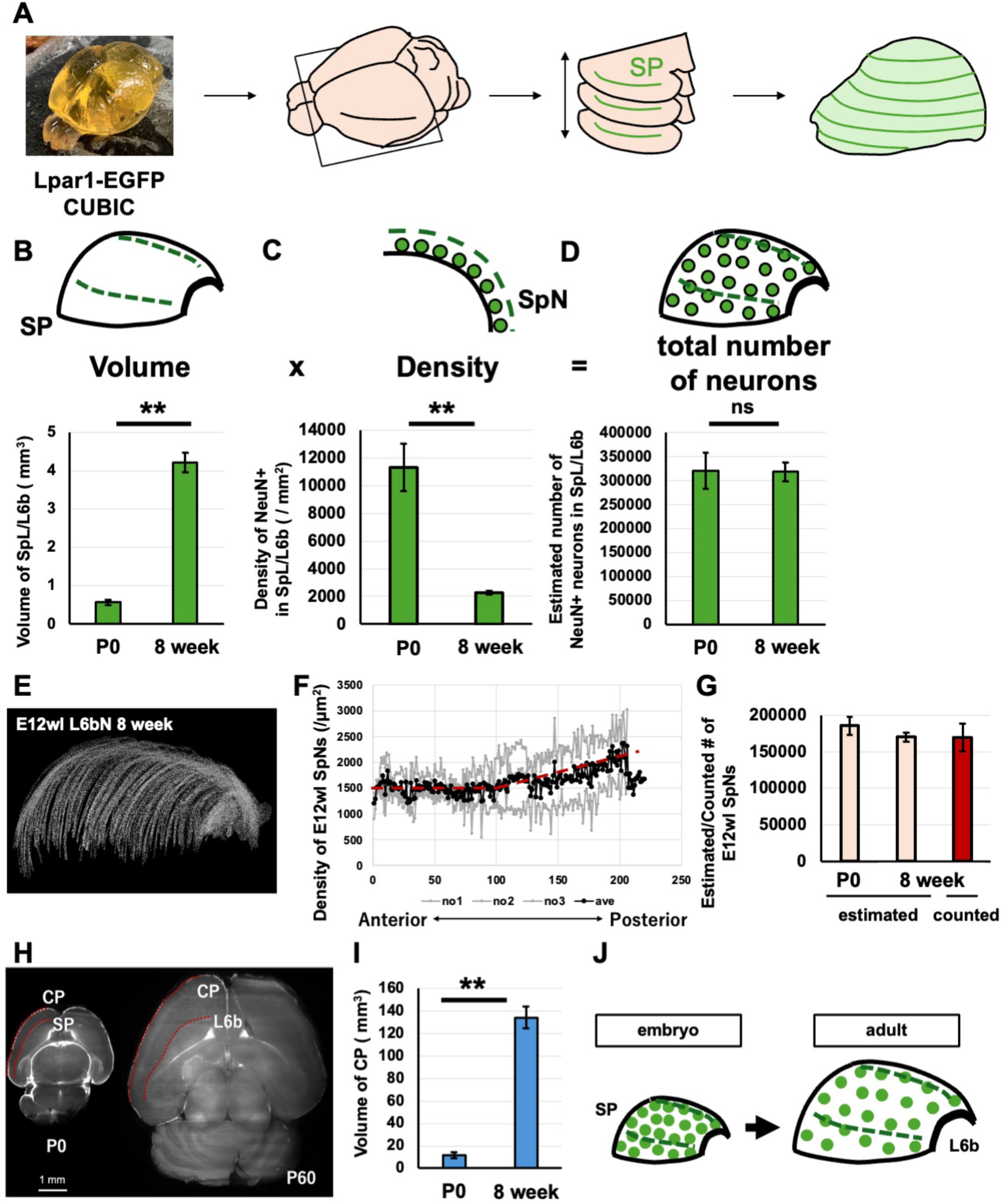
The total number of SpNs remains largely unchanged during postnatal development. (A) Schematic of the method used to measure the volume of the SP layer and L6b across the entire neocortex. The black outlined square indicates the frame of each image acquired by LSFM (left). Green lines mark the regions containing Lpar1-GFP+ cells in each acquired horizontal section (middle). The reconstructed 3D structure of the SP/L6b layer is shown on the right. (B) Three-dimensional reconstruction of the SP layer. Comparison of SP volume at P0 and L6b volume at 8 weeks (n = 4). (C) Schematic diagram of SpNs within cortical slice. Cell densities of NeuN+ cells in SP at P0 and L6b at 8 weeks were quantified from six fields of view per animal (n = 3 animals). (D) Schematic diagram of 3D reconstruction of the distribution of SpNs throughout the SP. Estimated total number of neurons in the SP at P0 and L6b at 8 weeks (n = 4). (E) The 3D-reconstructed half-hemisphere of sequential image of E12wl EdU+ L6b neurons (F) Density profile of E12wl EdU+ neurons in L6b along the anteroposterior (AP) axis. Gray lines represent individual data, and the black line shows the mean (n = 3 brains, 18 images). (G) Comparison of the actual counted number of EdU+ L6b neurons at 8 weeks with the estimated total number of SpNs at P0 and L6b neurons at 8 weeks from 3D reconstruction (n = 4 brains). (H) Horizontal sections of the whole brain of the Lpar1-GFP mouse in P0 (left) and 8 weeks (right). The red dashed line indicates the outline of the neocortex and the boundary between the neocortex and the SP layer/L6b. Scale bar = 1mm (I) Comparison of the volume of CP at P0 and 8 weeks (n = 4) (J) Schematic illustration of the postnatal expansion of the SP/L6b region. Green dots represent SpNs/L6b neurons. ∗P < 0.05, ∗∗P < 0.01, no significant (ns); Welch’s t-tests (unpaired two-tailed)

Next, we estimated the total number of SpNs at P0 and L6b neurons at 8 weeks in the entire brain. We calculated the densities of SpNs / L6b neurons based on NeuN+ cells at P0 (9148.73 ±2946.79 /μm^2^ n = 4) and at 8 weeks (2255.48 ±89.72/μm^2^, n = 4) (Fig 5C). By multiplying these neuronal densities by the respective SP and L6b volumes and dividing by the width of z-axis of slide section (20μm at P0 and 30 μm at 8 weeks), we estimated the total number of SpNs as 321,071 ±21,625, and the number of L6b neurons as 318,463 ±11,127 (Fig 3D). Based on this, the decrease in the SpN number was only 0.8%, with the loss 2,611 cells. Additionally, to determine whether this estimation effectively represented the actual number of SpNs / L6b neurons at each stage, we counted the number of E12wl L6b neurons in the entire neocortex regions by creating coronal section series slices. E12wl L6b neurons were distributed in the entire bottom part of the neocortex (Fig 5E). To evaluate the number of E12wl SpNs and L6b neurons, we performed the estimation in the same way as the above methods by using E12wl SpNs or L6Ns density data (Fig 5F). The estimated numbers of E12wl SpNs and L6b neurons were comparable to the counted values, showing no significant differences among them (Fig 5G). This result suggested that our estimation method accurately represent the actual number of L6b neurons.

Previous studies estimated the number of SpNs using the neocortical expansion ratio during postanal development. To determine whether the neocortex and SP/L6b expanded at different rates, we measured the volume of the neocortex using the same method at P0 and 8 weeks. The neocortex, defined as regions above SP/L6b and delineated by red lines (Fig 5H) increased from 11.60 ±1.40 mm^3^ at P0 (n = 4) to 134.35 ±5.67 mm^3^ at 8 weeks (n = 4, Fig 5I). This aligned with the volume measurement at respective stages by MRI (Zhang et al. 2005). Based on this, the volume expansion ratio of the neocortex was about 11.6 (P0/8 weeks) (Fig 5I). Considering that the expansion ratio of SP/L6b was about six times, these results indicate that the neocortex expands larger than SP/L6b. Moreover, the results imply that previous studies used a higher expansion ratio as SP ratio than actual and overestimated the number of SP neurons in the adult. Therefore, these findings indicate that the decrease in SpN density is primarily attributable to the expansion of the neocortex during postnatal development rather than to significant cell loss (Fig 5J).

### Layer-specific estimation of neuron numbers efficiently reflects a decrease

We next evaluated whether this estimation method accurately reflects the actual reduction in neuron numbers when neurons are eliminated. We focused on another early-born neuron population, CR cells, as CR cells are known to decrease through apoptosis during postnatal development (Anstotz et al. 2014; Ledonne et al. 2016; Zecevic and Rakic 2001) Indeed, the number of EdU+ cells in L1 was reduced at P14 (Fig 1G) and 8 weeks (Fig 6A). Using the same tissue clearing of the whole brain, we measured the volume of the MZ/L1 region (Fig 6B). The volume of MZ/L1 was 1.91 ±0.37 mm^3^ (n = 4) at P0 and 14.62 ±1.91 mm^3^ (n = 4) at 8 weeks (Fig 6C), indicating approximately a 7-fold increase. We next analyzed the density of NeuN+ in MZ/L1 at P0 and 8 weeks. The density was 2,650.16 /mm^2^ at P0 and 328.46 /mm^2^ at 8 weeks, which showed a 6.5-fold decrease (Fig 6D). We multiplied the volume and density to estimate the number of MZ/L1 neurons. The number was 253,357 at P0 and 160,017 at 8 weeks (Fig 6E), indicating that the number of MZ/L1 neurons decreased by about 40%. Since neurons were defined based on the expression of NeuN, the number of neurons in MZ/L1 may include L1 interneurons (Schuman et al. 2019). However, since not only CR cells but also interneurons decrease during postnatal development, the 40% decrease in MZ/L1 neurons was reasonable. These results indicate that our estimation method successfully captures the actual reduction in the total neuron numbers. Furthermore, this approach enables to analyze the dynamics of neuron number without the confounding effects of neocortical expansion during postnatal development.

**Fig 6.**
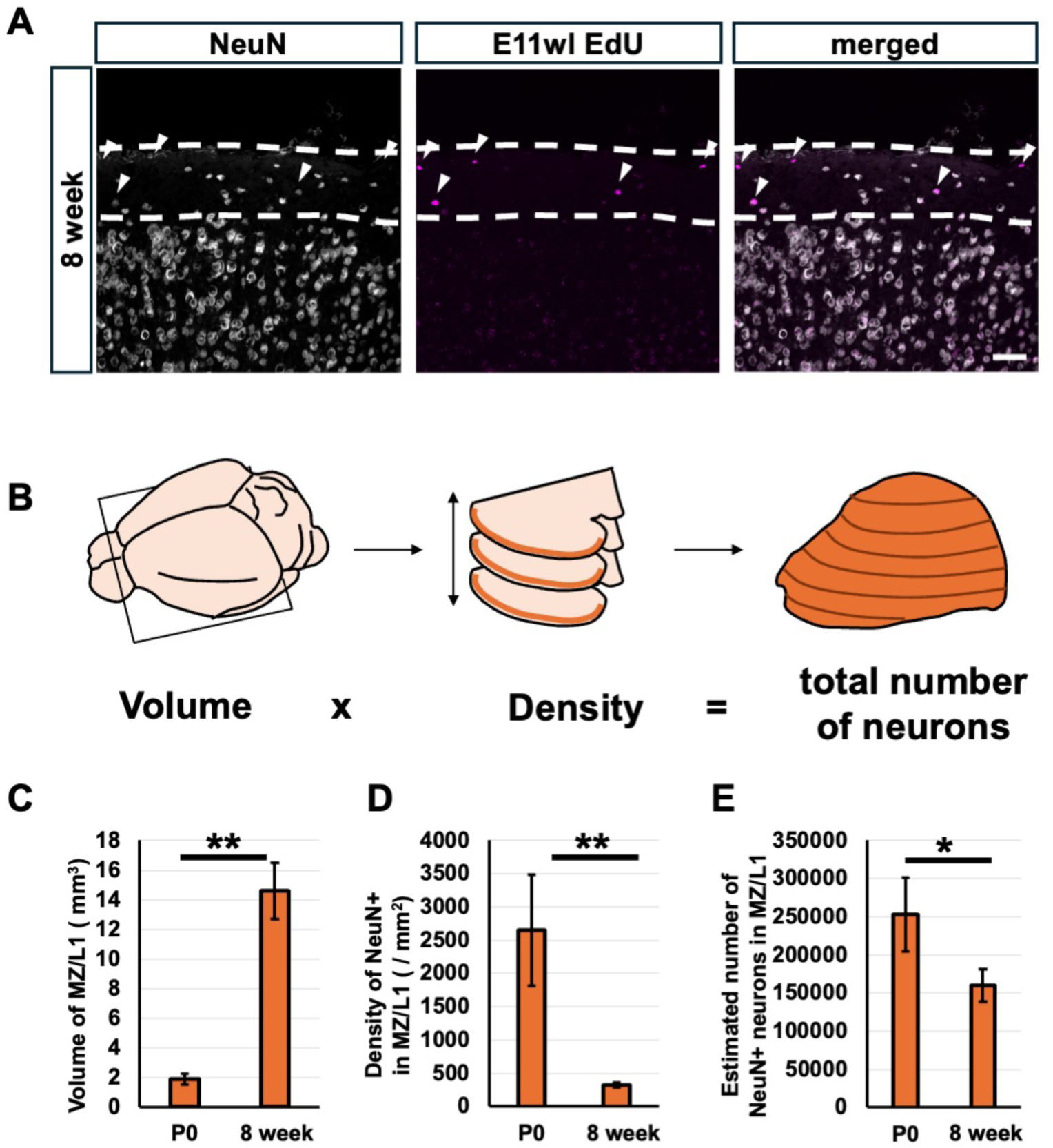
The decrease in Cajal–Retzius (CR) cells is detected by the 3D estimation method of cell number. (A) Immunohistochemical images showing the distribution of E11 EdU+ cells in layer1(L1) in the adult. NeuN (white) and EdU (red) staining at 8 weeks. White arrowheads show EdU+ cells. White dashed lines delineate the boundary of L1. Scale bar = 50μm (B) Schematic of the method used to measure the volume of the marginal zone (MZ)/L1 across the entire neocortex. The black-outlined square indicates the frame of each image acquired by LSFM (left). Orange bands indicate the MZ/L1 regions in each horizontal section (middle). The reconstructed 3D structure of the entire MZ/L1 layer is shown on the right. (C) Comparison of the volume of the MZ layer at P0 and L1 at 8 weeks (n = 4). (D) Density of NeuN+ cells in the MZ at P0 and L1 at 8 weeks (/mm^2^) (n = 6 field of views, 3 animals) (E) Estimated total number of neurons located in the MZ at P0 and L1 at 8 weeks (n = 4) ∗∗P < 0.01, no significant (ns); Welch’s t-tests (unpaired two-tailed)

## Discussion

SpN are essential for embryonic neocortical development, yet they have long been regarded as a largely transient population that undergoes substantial postnatal elimination in neocortex (Allendoerfer and Shatz 1994). In this study, by combining extended-window birthdate labeling with whole brain 3D quantification, we demonstrate that the majority of early-born SpNs persist into adulthood and constitute the core neuronal population of layer 6b(L6b) in the mouse neocortex. Our findings reveal that the developmental decline in SpN density is not caused by widespread programmed cell death but is instead explained by tangential expansion of the cortical mantle during postnatal growth. While some studies have suggested the connection between SP and L6b neurons in rodents (deFazio et al. 1987; Pedraza et al. 2014), definitive evidence had been lacking. Our study provides quantitative evidence that SpNs do not disappear after birth rather persist as L6b neurons, thereby revising a long-standing view of the developmental fate of the Subplate.

### Revisiting the long-standing assumption of postnatal SpN loss

Previous birthdating studies (Allendoerfer and Shatz 1994; Hoerder-Suabedissen and Molnar 2013; Price et al. 1997) included that most SpNs undergo extensive postnatal cell death. However, these studies relied on single-pulse labeling of thymidine analogs, which labels only a small subset of cells born within a narrow 2–6 h window. In addition, they quantified SpNs only in small tissue sections without measuring the total number of neurons or the regional volume. Because the neocortex undergoes substantial volumetric expansion during early postnatal development (Heumann et al. 1978; Kronman et al. 2024), density-based measurements inevitably give the impression of cell loss even when absolute cell number is preserved. Indeed, our extended-window EdU-labeling increased labeling efficiency 5-6 fold compared with single injections‘Fig1C,F’, and our volumetric data showed a 7.4-fold increase in SP/L6b volume from P0 to adulthood (Fig.5B). These methodological advances directly address the limitations of earlier studies and indicate that much of the reported decline in SpN density can be explained by postnatal cortical expansion rather than widespread programmed neuronal elimination. Notably, although the SP/L6b expanded by approximately 7.4-fold from P0 to adulthood, the neocortex as a whole expanded even more by about 11-fold (Fig.5I).

Because density inevitably decreases as tissue volume increases, this difference in expansion rates explains much of the apparent decline in SpN density, without requiring large-scale programmed neuronal loss. This also implies that earlier density-based studies, which did not account for differential expansion of SP versus the whole neocortex, may have overestimated the extent of postnatal SpN loss.

### Extended-window birthdating captures the majority of SpNs

By administrating EdU at multiple time points across the full neurogenic window of SpNs(E10.5-E12.5), we achieved comprehensive labeling of early-born neurons that was not possible in earlier studies using single-pulse BrdU. The EdU^+^ fraction within L6b neurons was markedly higher than previously reported, consistent with the extended labeling window capturing both direct ventricular zone-derived neurons and a substantial population of intermediate progenitor-derived SpNs. More than 90 % of L6b neurons were labeled by our E10-12 whole-labeling protocol, indicating that the vast majority of L6b neurons originated from early-born SpNs and survive into adulthood. A slightly higher-than-expected EdU⁺ ratio in the E12wl group may be explained by the contribution of intermediate progenitor–derived SpNs, consistent with previous evidence that ∼40% of SpNs originate from IP cells (Vasistha et al. 2015).

### Whole-brain quantification reveals minimal postnatal SpN elimination

The combination of CUBIC tissue clearing and light-sheet fluorescence microscopy enabled full reconstruction of SP/L6b region and direct estimation of total neuron number. Despite a 7.4-fold increase in volume, the total number of SpNs changed by <1% between P0 and adulthood, demonstrating minimal developmental elimination. Importantly, our estimation method successfully detected the well-established ∼40% reduction of Cajal-Retzius (CR) cells, confirming that the approach is sensitive to genuine neuronal loss. These results validate our conclusion that SpNs- and not CR cells-exhibit remarkable postnatal persistence. Although regional heterogeneity cannot be fully excluded, we minimized sampling bias by collecting multiple fields from matched anatomical levels and averaging across them.

### Molecular heterogeneity and developmental origins of surviving SpNs

We further identified molecular differences among surviving SpNs based on birthdate. E12-born SpNs exhibited higher expression of Nurr1 and Tbr1 compared to E11-born neurons, consistent with earlier observations of SpN heterogeneity. This birthdate-dependent heterogeneity is likely linked to distinct developmental origins: early-born SpNs (E11) include neurons migrating tangentially from the rostromedial telencephalic wall (RMTW), while later-born SpNs (E12) arise mainly from the ventricular zone (Pedraza et al. 2014). In addition, a subset of SpNs has been reported to originate from the dorsomedial cortical hem (Saito et al. 2019), further supporting the presence of multiple birthdate- and origin-dependent subtypes. RMTW-derived neurons are generated around E11 and migrate tangentially into the subplate, whereas later-born SpNs (E12) mainly arise from the ventricular zone (Pedraza et al. 2014).

These developmental origins are consistent with our finding that E11wl and E12wl SpNs show distinct molecular profiles, suggesting that early-born SpNs comprise multiple birthdate- and origin-dependent subtypes. Thus, the adult L6b contains multiple molecularly and developmentally defined subtypes of SpNs, raising the possibility that these subtypes contribute differentially to adult cortical circuit.

### Functional implications for adult L6b circuits

Although the precise roles of L6b neurons in adult cortical processing remain to be fully clarified, accumulating evidence suggests that they are functionally active and contribute to sensory and higher-order computations (Kim et al. 2025; Messore et al. 2025; Zolnik et al. 2024). CTGF-expressing L6b neurons exhibit state-dependent activity and long-range projections to higher-order thalamic nuclei, while entorhinal L6b neurons contribute to hippocampal-driven spatial memory (Ben-Simon et al. 2022; Kim et al. 2025; Zolnik et al. 2020).

Recent studies have further demonstrated that L6b neurons actively participate in functional circuits with sensory modality–specific roles. In the visual cortex, CTGF-expressing L6b neurons exhibit experience-dependent activity and plasticity (Yoneda et al. 2023), while other studies have reported L6b contributions to both visual and auditory processing (Chang et al. 2024). In our study, we found that early-born, EdU⁺ L6b neurons include CTGF⁺ cells, suggesting that surviving SpNs may likewise integrate into mature functional circuits in the adult neocortex. Elucidating how embryonic SpNs transition into these functional adult L6b neurons—both molecularly and electrophysiologically—will be an important direction for future research. Our finding that most CTGF^+^ neurons are early-born SpNs strongly suggests that SpNs, long viewed as exclusively embryonic players, continue to participate in mature neocortical circuits. Understanding how these neurons transition from transient embryonic organizers to stable adult circuit components will be an important focus for future studies.

### Species difference in subplate development

While rodents retain the majority of SpNs into adulthood, primates-including humans-show dramatic expansion and subsequent reorganization of the subplate during development (Adorjan et al. 2019; Huang et al. 2009; Judas et al. 2013; Kanold 2009; Kostovic et al. 2011; Kostovic et al. 2019; Kostovic 2020). In species with large and protracted neurogenesis, the subplate forms a massive transient compartment enriched in ECM and diverse transient neuronal populations. In these species, genuine developmental elimination of SpNs is more pronounced and may contribute to emergence of specialized thalamocortical pathways and complex cognitive functions. Thus, SpN fate is likely species-specific, and our results define the rodent baseline against which primate-specific evolutionary adaptations can be interpreted.

### Conclusion

Together, our findings overturn the long-standing view that the rodent subplate is largely eliminated after birth. Instead, early-born SpNs survive, persist as L6b neurons, and likely contribute to mature cortical computation. By integrating extended window birthdate labeling with whole-brain quantitative imaging, this study reconciles decades of conflicting observations and establishes a revised framework in which the subplate is not merely a transient embryonic scaffold but the developmental origin of persistent and functionally important layer of adult cortex.

## Acknowledgments

We thank M. Sato (Kanazawa University) for the valuable discussion about the estimation of SpN number, T. Kumamoto (Tokyo Metropolitan institute of Medical Science) for teaching us how to use CUBIC clear tissue method, and S. Sakai (Institute of Science Tokyo) for teaching us how to use Light Sheet fluorescence microscopy. We also thank members of the Neural Development Project of Tokyo Metropol. Inst. Med. Sci. and Development Biology Lab of Waseda Univ. for discussion and comments on the study. Y.S. was supported in part by Waseda University Early Bird Program.

## Author contributions

Author contributions: Y.S., K.M-I.,a nd C.O-M. designed research; Y.S., and K.M-I., performed research; Y.S., K.M-I, and C.O-M., analyzed data; Y.S., K.M-I, C.H., and C.O-M., wrote the paper; C.H., imbued advice and expertise, and contributed to the manuscript; C.O-M provided financial and administrative support, supervise, and contributed to the manuscript.

---

**Supplement figure 1.**
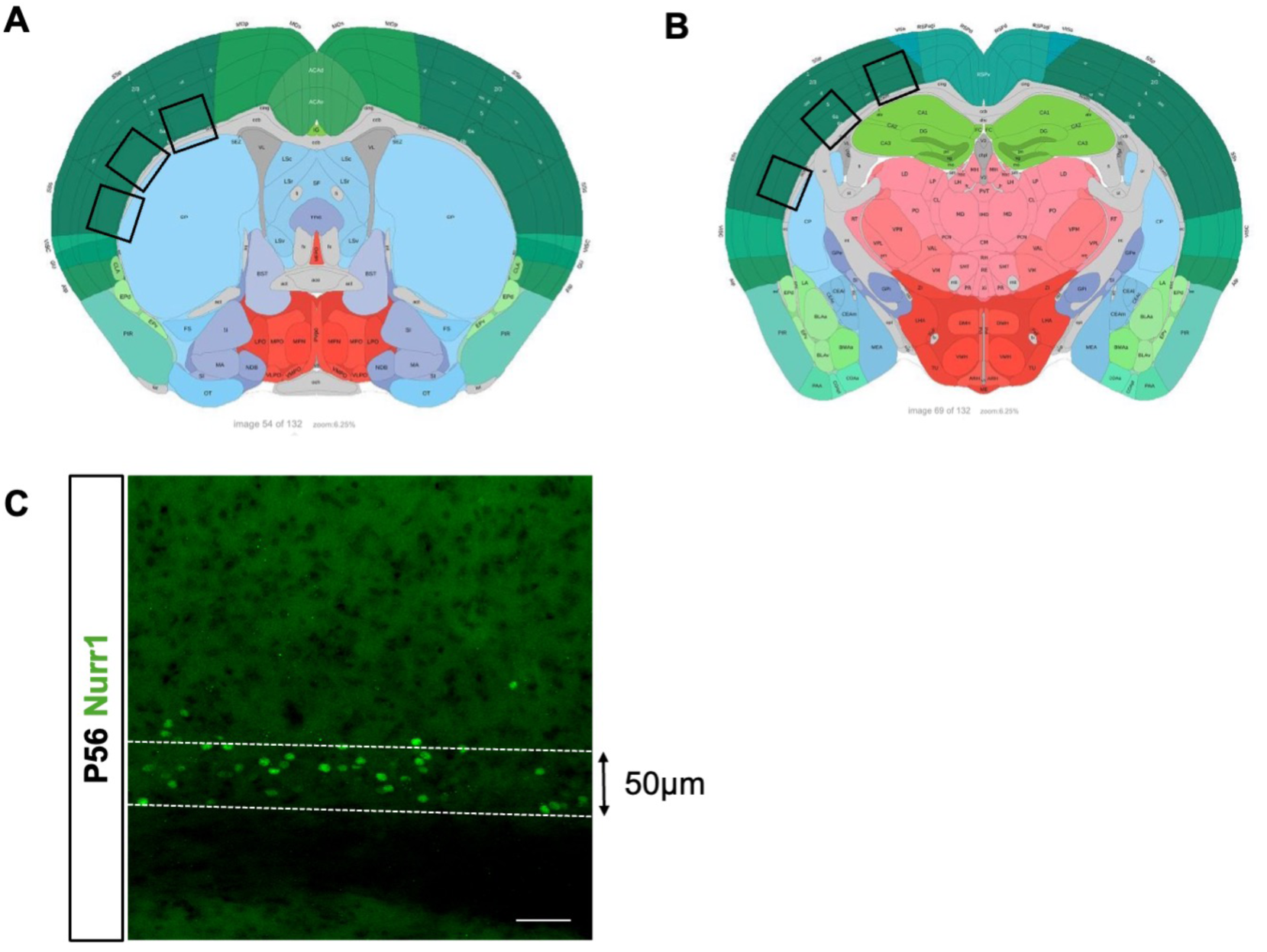
Window view of images for quantification and definition of L6b. (A, B) Serial coronal sections from the mouse brain atlas from Allen brain atlas No.54 (A) or No.69 (B). Black squares indicate the imaging area used for quantification (a total of 6 fields). (C) Definition of the SP layer/L6b. The boundary between SP layer/L6b and the neocortex was defined as 50μm above the white matter (WM).

**Supplement figure 2.**
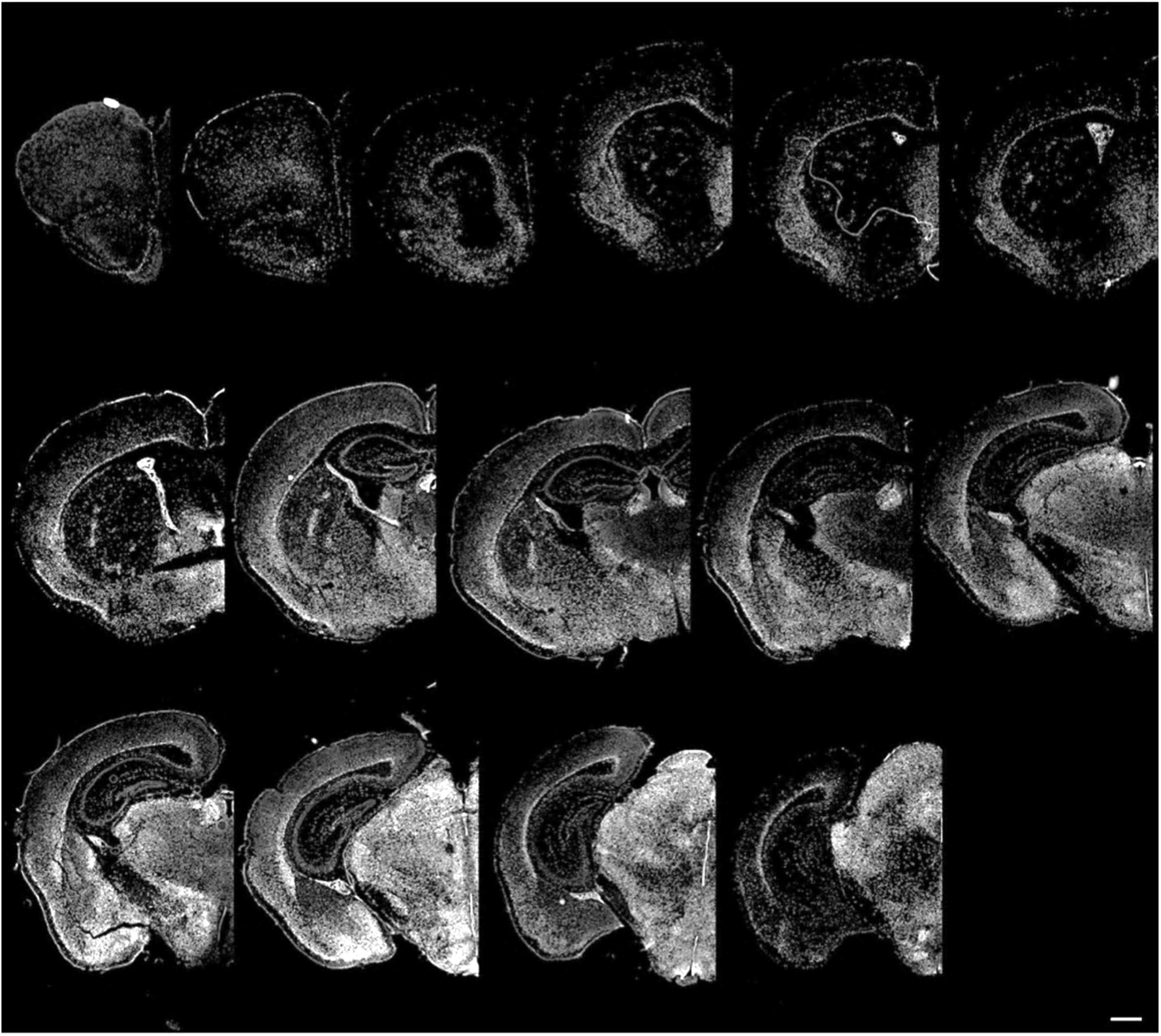
Serial coronal sections of an E11wl mouse brain. EdU positive cells represent neurons born within the E11 labeling window. Scale bars indicate 500μm.

**Supplement figure 3.**
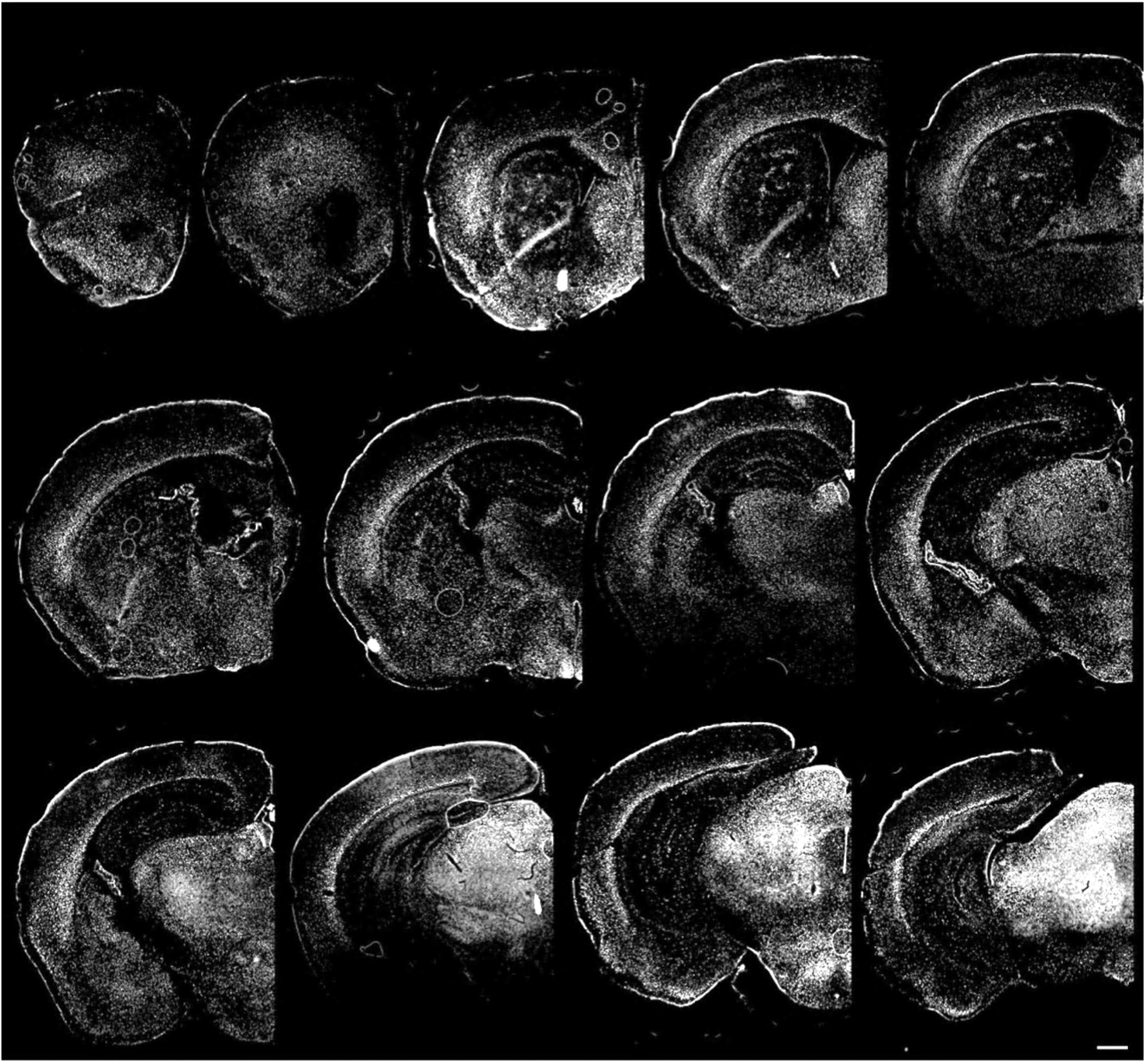
Serial coronal sections of an E12wl mouse brain. EdU positive cells represent neurons born within the E12 labeling window. Scale bars indicate 500μm.

**Supplement figure 4.**
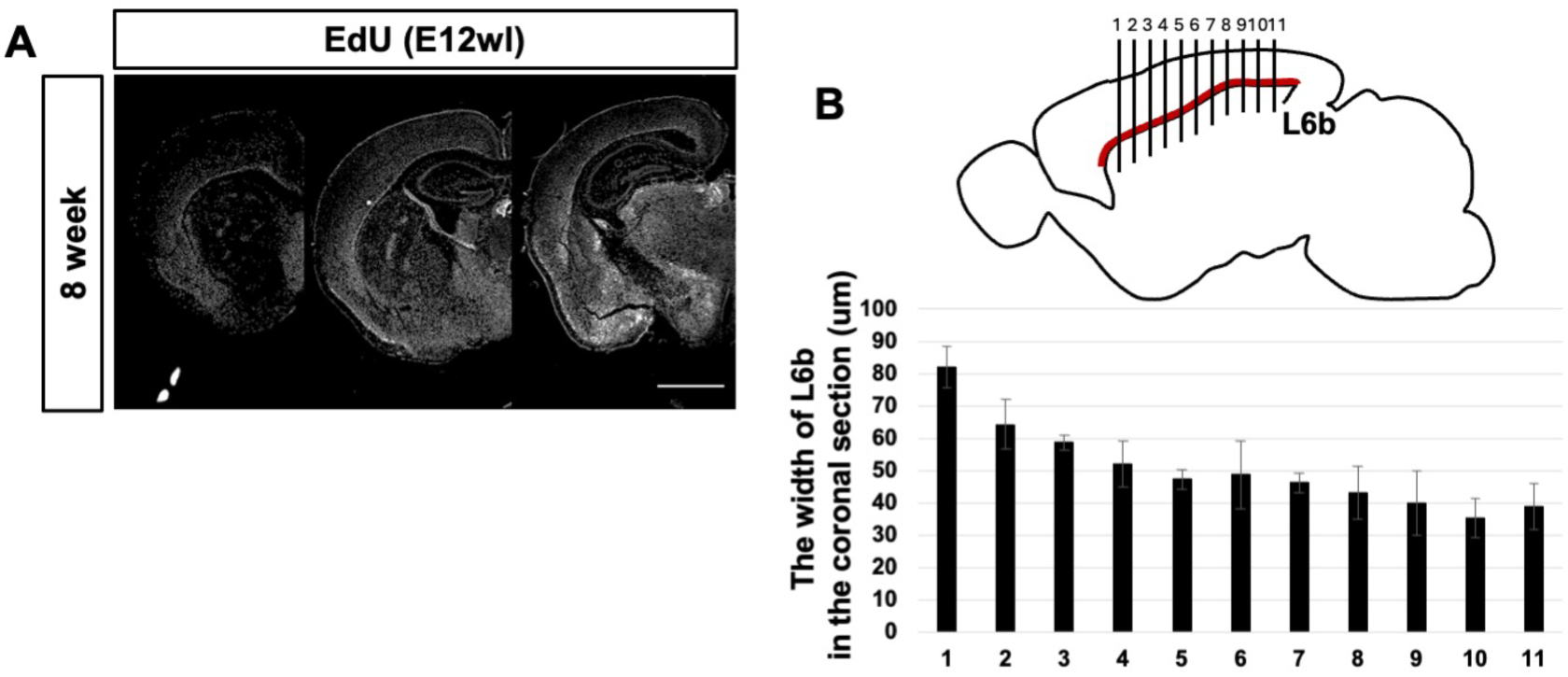
Distribution of EdU+ cells across coronal sections and thickness of layer 6b neurons (L6bNs). (A) Series of coronal sections of confocal micrographs of E12wl EdU+ cells in 8 weeks, which were rostral, middle, or caudal. (B) The width of L6b in 8 weeks in the coronal section (μm). The horizontal axis indicates the position along the rostrocaudal axis, measured as the distance from the junction between the olfactory bulb and neocortex (μm). The average width of L6b across all sections was 50.60 μm.

**Supplement figure 5.**
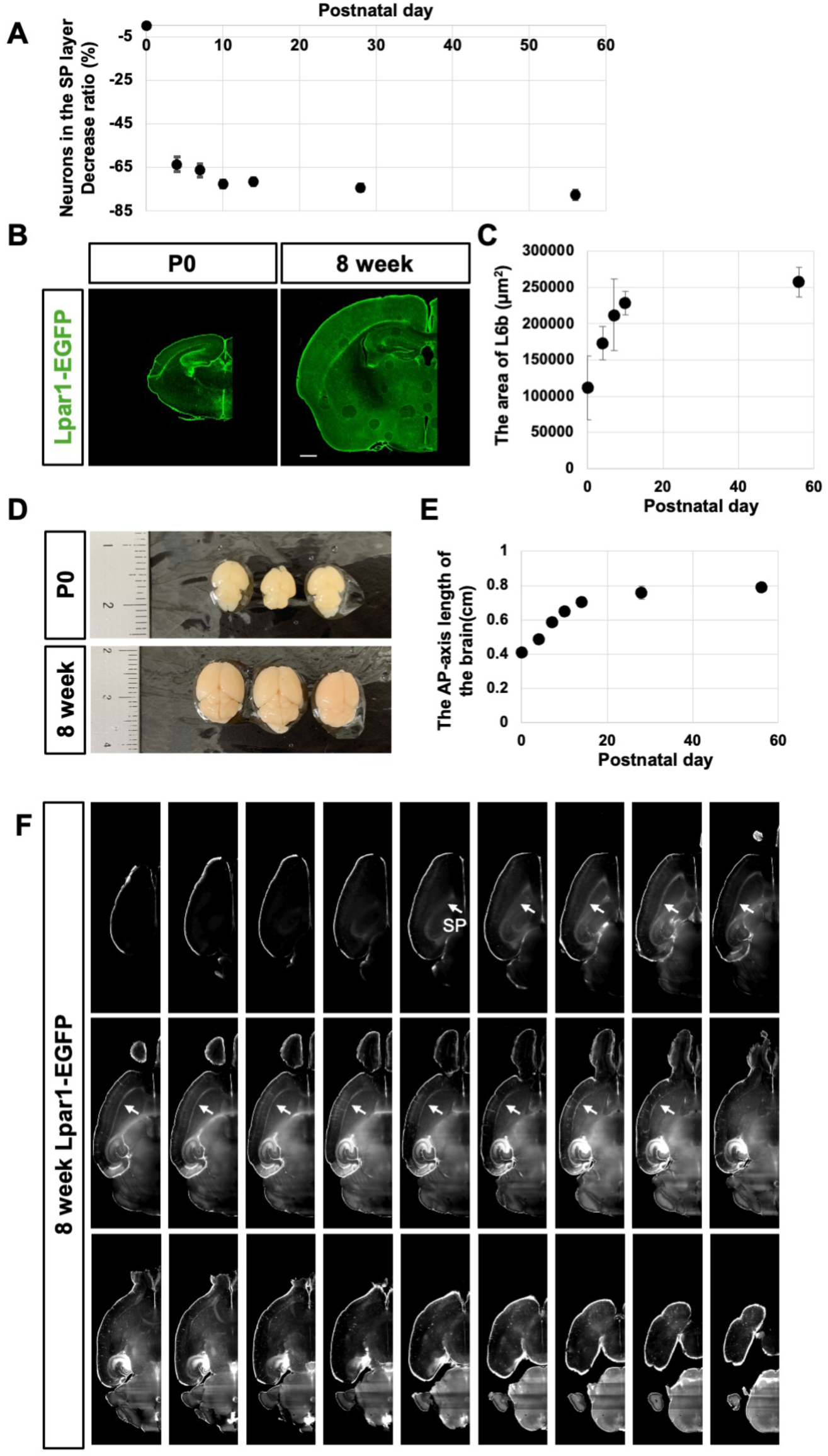
Postnatal volume expansion of the mouse cortex. (A) Decrease rate of the neuronal (NeuN+) density in the SP during postnatal development. (The density in each stage – in P0)/(the density in P0) x100. (n = 3 brains, n = 18 images) (B) Immunohistochemical images of the coronal sections of Lapr1-GFP mouse brain at P0 (left) and 8 weeks (right) (C) Area of SP/L6b is defined by Lpar1-GFP expression (μm^2^) (n=3). (D) Photographs of 4%PFA fixed mouse brain in P0 (top) and 8 weeks (bottom) (E) Anteroposterior (AP) length of the brain excluding the cerebellum (cm) (n=3) (F) Series of horizontal sections of 8-weel old Lpar1-GFP mouse brain, imaged by Light sheet fluorescence microscopy. Sections were taken at 200μm intervals. White arrows indicate the Lpar1-GFP+ SP layer.

